## Supplemental data for "Metabolic and neurobehavioral disturbances induced by purine recycling deficiency in *Drosophila*"

#### Supplementary Files

- **Supplementary figures**
-
- **Supplementary tables**
-
- **Supplementary movies**

#### Supplementary figures

**A**

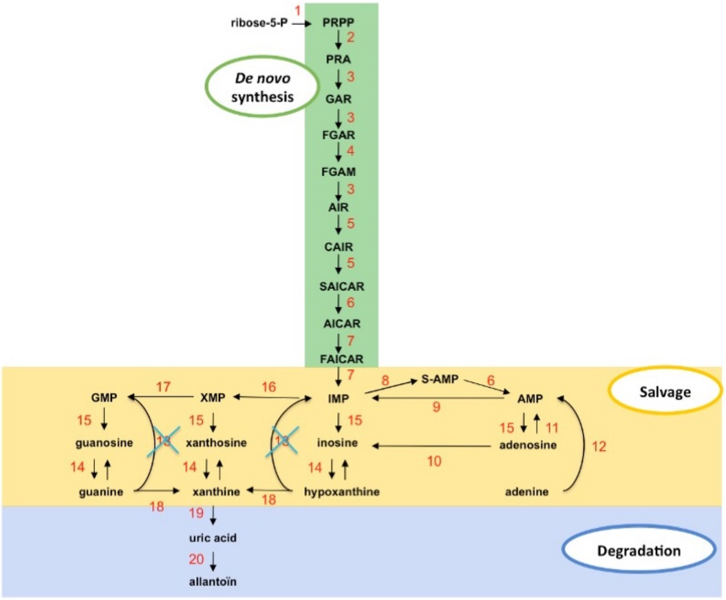


**B**

| Step | Enzyme | Human | *Drosophila* | % Similarity | % Identity |
|---|---|---|---|---|---|
| 1. | PRPS | *PRPS1* | *CG6767* | 94 | 88 |
| 2. | GPAT | *PPAT* | *Prat* | 74 | 61 |
| 2. | GPAT | *PPAT* | *Prat2* | 75 | 61 |
| 3. | GARS/AIRS/GART | *GART* | *Gart/ade3* | 65 | 48 |
| 4. | FGAMS | *PFAS* | *Pfas/ade2* | 64 | 50 |
| 5. | AIRC/SAICARS | *PAICS* | *Paics/ade3* | 74 | 59 |
| 6. | ADSL | *ADSL* | *AdSL* | 80 | 68 |
| 7. | ATIC | *ATIC* | *CG11089* | 83 | 71 |
| 8. | ADSS | *ADSS2* | *AdSS* | 76 | 63 |
| 9. | AMPD | *AMPD1* | *AMPdeam* | 71 | 54 |
| 10. | ADA | *ADA* | *Ada* | 41 | 25 |
| 11. | ADK | *ADK* | *AdenoK* | 68 | 52 |
| 12. | APRT | *APRT* | *Aprt* | 65 | 46 |
| 13. | HGPRT | *HPRT1* | / | / | / |
| 14. | PNP | *PNP* | *CG16758* | 68 | 51 |
| 15. | 5-NT | *NT5C2* | *CG32549* | 72 | 60 |
| 16. | IMPDH | *IMPDH1* | *raspberry* | 83 | 69 |
| 17. | GMPS | *GMPS* | *burgundy* | 76 | 59 |
| 18. | GDA | *GDA* | *DhpD* | 58 | 43 |
| 19. | XDH/XO | *XDH* | *rosy* | 68 | 52 |
| 20. | UO | / | *Uro* | / | / |

**Figure S1.** Comparison of purine metabolism pathways in *Drosophila* and humans. (**A**) Schematic diagram of the purine biosynthesis and degradation pathways in *Drosophila melanogaster*, based on sequence homology with the human genes. (**B**) Percent sequence similarity between human and *Drosophila* homologs of purine metabolism enzymes. Note the lack of urate oxidase (UO, *step 20*) in humans, due to a primate-specific loss of this gene, and the fact that *Drosophila* does not have a homolog of human HGPRT (*step 13*). However, the APRT enzyme has been conserved (*step 12*), suggesting that it is the only recycling enzyme of the purine salvage pathway in *Drosophila*.

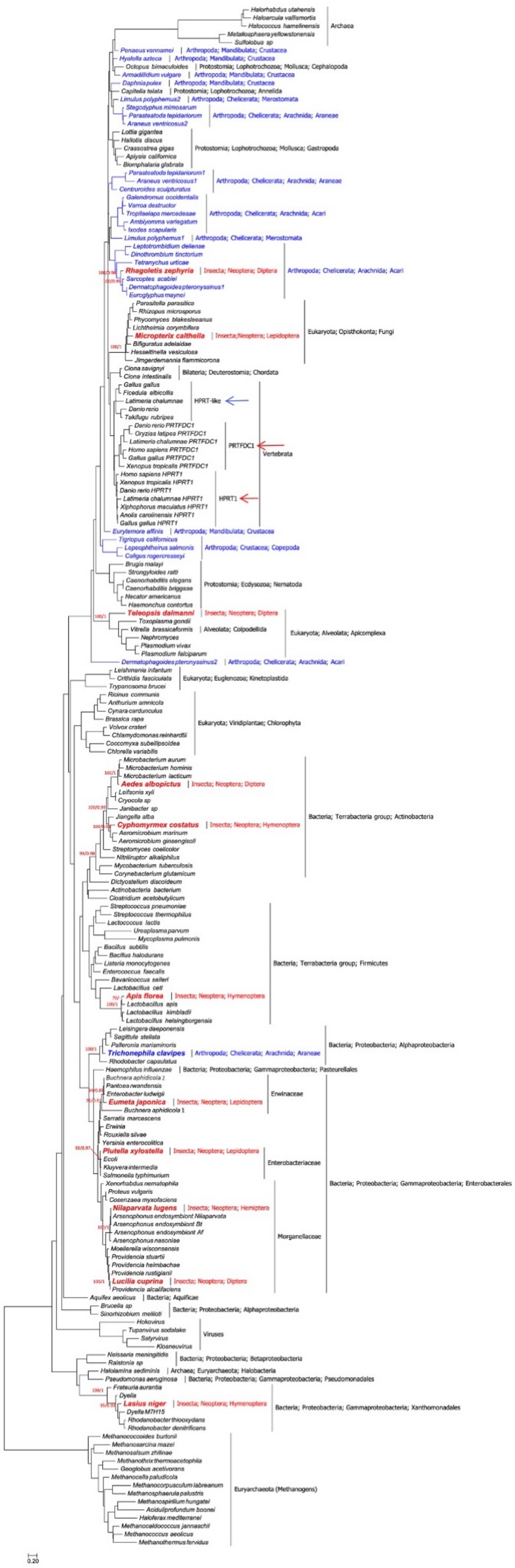
Archaea
Arthropoda; Mandibulata; Crustacea
Protostomia; Lophotrochozoa; Mollusca; Cephalopoda
Protostomia; Lophotrochozoa; Annelida
Arthropoda; Chelicerata; Merostomata
Arthropoda; Chelicerata; Arachnida; Araneae
Protostomia; Lophotrochozoa; Mollusca; Gastropoda
Arthropoda; Chelicerata; Arachnida; Acari
Insecta; Neoptera; Diptera
Eukaryota; Opisthokonta; Fungi
Insecta;Neoptera; Lepidoptera
Bilateria; Deuterostomia; Chordata
HPRT-like
PRTFDC1
Vertebrata
HPRT1
Arthropoda; Crustacea; Copepoda
Protostomia; Ecdysozoa; Nematoda
Alveolata; Colpodellida
Eukaryota; Alveolata; Apicomplexa
Eukaryota; Euglenozoa; Kinetoplastida
Eukaryota; Viridiplantae; Chlorophyta
Bacteria; Terrabacteria group; Actinobacteria
Insecta; Neoptera; Hymenoptera
Bacteria; Terrabacteria group; Firmicutes
Bacteria; Proteobacteria; Alphaproteobacteria
Bacteria; Proteobacteria; Gammaproteobacteria; Pasteurellales
Erwinaceae
Insecta; Neoptera; Lepidoptera
Enterobacteriaceae
Bacteria; Proteobacteria; Gammaproteobacteria; Enterobacterales
Insecta; Neoptera; Hemiptera
Morganellaceae
Bacteria; Aquificae
Viruses
Bacteria; Proteobacteria; Betaproteobacteria
Archaea; Euryarchaeota; Halobacteria
Bacteria; Proteobacteria; Gammaproteobacteria; Pseudomonadales
Bacteria; Proteobacteria; Gammaproteobacteria; Xanthomonadales
Euryarchaeota (Methanogens)
0.20

**Figure S2.** Unrooted Maximum Likelihood phylogeny of HGPRT proteins (189 taxa, 130 sites). Phylogenetic analyses show that: (a) HGPRT proteins are ancient, as they are present in Bacteria and Archaea. (b) Two paralogs (HPRT1 and PRTFDC1) are found in human and vertebrates (*red arrows*). An additional protein (HGPRT-like) is found in other vertebrates (*blue arrow*). (c) Of special interest for our study is that HGPRT proteins are very rare in Insects and, in particular, are absent in Drosophilidae, with one possible exception (see main text and Figure S3). Indeed, this preliminary phylogenetic analysis shows that insect HGPRT proteins cluster mainly with Bacteria (but also with Fungi, Apicomplexa and Acari). This strongly suggests that all the 11 HGPRT proteins found to date in Insecta (*in bold red font*) were very likely acquired by horizontal gene transfer. (d) The potential horizontal gene transfer event in the spider *Trichonephila clavipes*, which clusters with Alphaprotoebacteria, is highlighted in *bold blue font*. All other Arthropod sequences are shown in *blue font*. Branch-support values, only given at relevant nodes, are UFBoot/Bayesian Posterior Probabilities. The scale bar indicates the estimated number of substitutions per site.

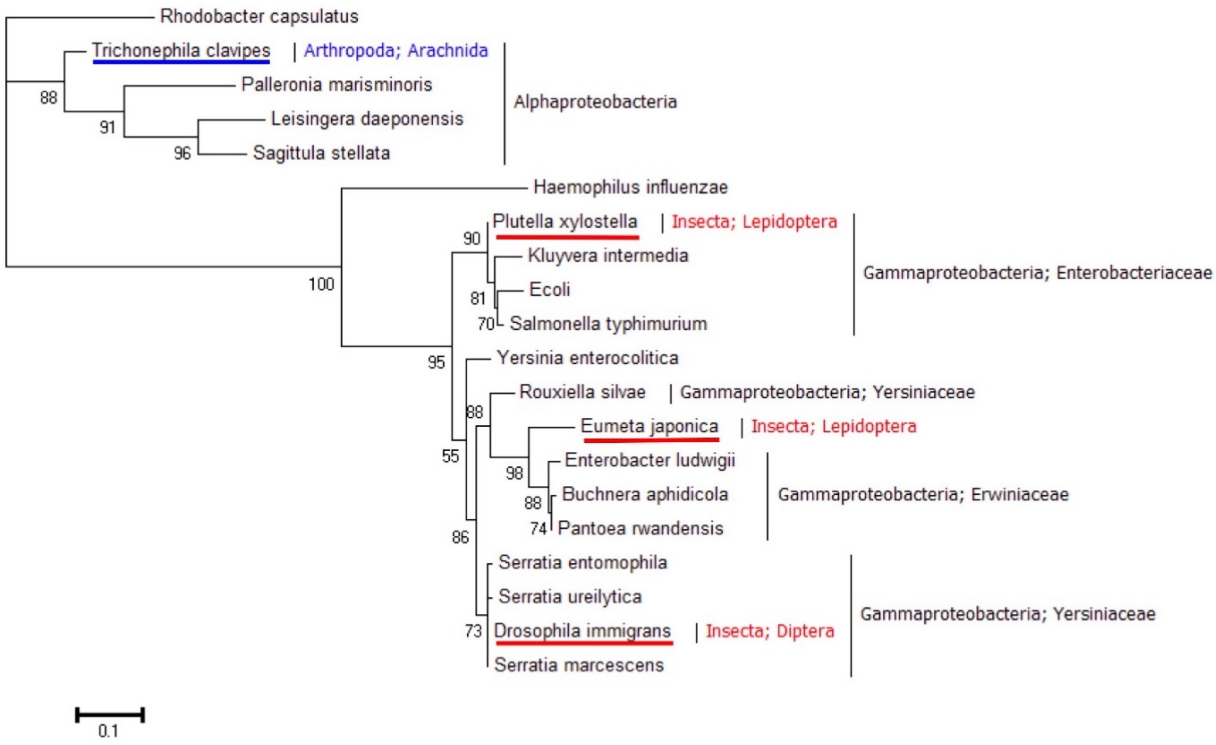


**Figure S3.** Unrooted maximum likelihood phylogeny of HGPRT proteins (20 taxa, 177 sites). The dataset comprises 17 relevant taxa used in Figure S2 together with the HGPRT proteins of *D. immigrans*, *Serratia ureilytica* and *Serratia entomophila.* The *Drosophila immigrans* sequence (*red underline*) clusters with the gammaproteobacterial genus *Serratia*. This can be interpreted either a contamination of the sequenced sample (*D. immigrans* and *S. marcescens* proteins are 100% identical) or a very recent horizontal gene transfer event. The potential horizontal gene transfer events in *Plutella xylostella* and *Eumata japonica* (also shown in Figure S2) are also highlighted (*red underline*) as well as that of the arachnid *Trichonephila clavipes* (*blue underline*). Branch-support values are aLRT (SH-like). The scale-bar indicates the estimated number of substitutions per site.

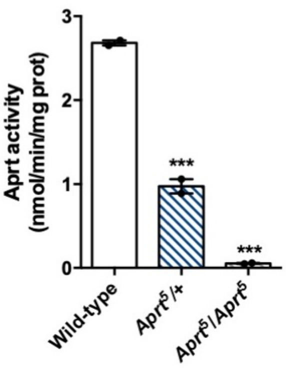


**Figure S4.** Lack of Aprt enzymatic activity in the *Aprt*$^5$ mutant. Aprt activity was assayed on whole adult fly extracts, showing that it is strongly reduced in heterozygous *Aprt*$^5$/+ mutants, and absent in homozygous *Aprt*$^5$ flies, compared to the wild-type. The fact that it is decreased more than 2-fold in heterozygous flies may suggest a dominant negative effect of the mutation. 2 independent determinations were performed on 20 whole flies per genotype. One-way ANOVA with Dunnett's *post-hoc* test for multiple comparisons (***$p < 0.001$).

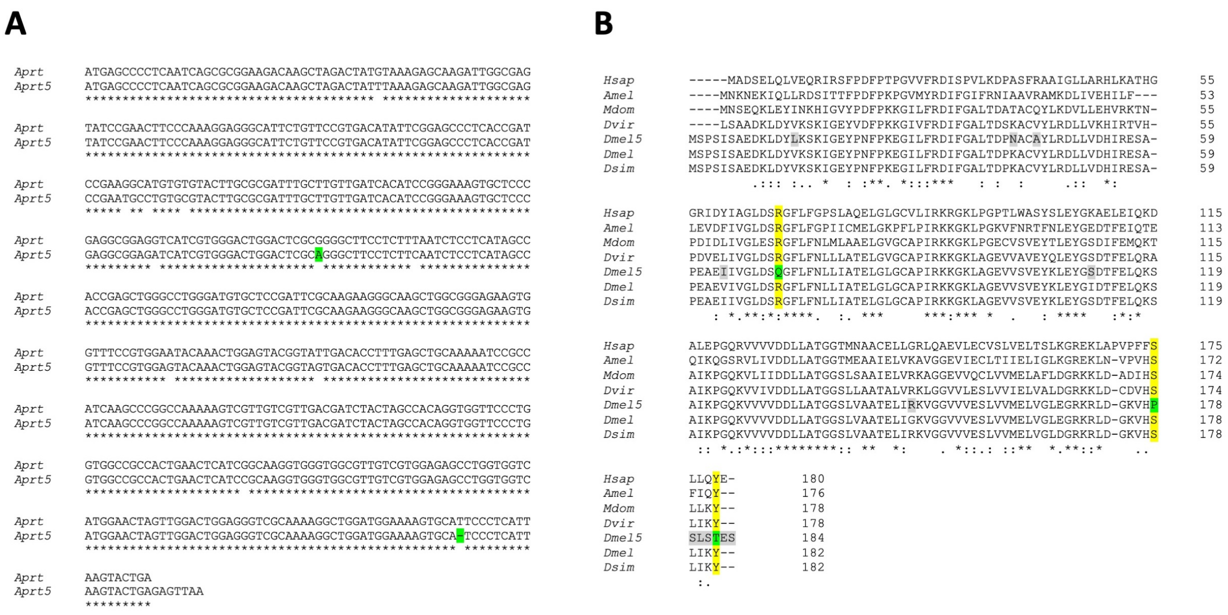


**Figure S5.** Alignment of wild-type and mutant *Aprt* cDNAs and predicted protein sequences. (**A**) Nucleotide sequence alignement of the coding regions in *Drosophila Aprt* and *Aprt*$^{5}$ cDNAs. A base substition and a deletion that are responsible for the three prominent mutations in the Aprt5 protein (see in B) are highlighted in green. (**B**) Alignment of human and insect Aprt proteins. The residues modified in *Drosophila* Aprt5 (Dmel5) compared to *Drosophila* wild-type Aprt (Dmel), are highlighted in gray when they correspond to a variable residue in different sequences, and in green when they alter amino acid residues that were highly conserved throughout evolution. The R71Q, S178P and Y182T mutations change three residues (highlighted in yellow) that were conserved in Aprt sequences from *Drosophila* to humans. These mutations are therefore likely to be responsible for the loss of enzymatic activity in Aprt5. *Hsap*: *Homo sapiens*, *Amel*: *Apis mellifica*, *Mdom*: *Musca domestica*, *Dvir*: *Drosophila virilis*, *Dmel5*: *Drosophila melanogaster* Aprt5, *Dmel: Drosophila melanogaster* wild-type Aprt, *Dsim*: *Drosophila simulans*. The nucleotide and amino acid sequences were retrieved from FlyBase and GenBank, except for *Drosophila* mutant Aprt5, which was sequenced in this work. The alignements were performed using MUSCLE (https://www.ebi.ac.uk/Tools/msa/muscle/) and Clustal Omega (https://www.ebi.ac.uk/Tools/msa/clustalo/) multiple sequence alignment tools.

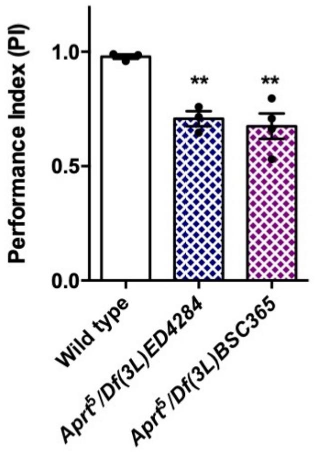


**Figure S6.** SING behavior of hemizygous *Aprt* mutant flies. *Df(3L)ED4284 and Df(3L)BSC365* are deficiencies located in 62B4-B12 and 62B7-D3, respectively, that removes *Aprt* and several neighbor genes. Hemizygous *Aprt*$^5$ flies, in which the mutation was placed over these deficiencies, showed an early locomotor decline in the SING assay at 10 d a.E., similarly to homozygous *Aprt*$^5$ flies. Results of 3-4 independent experiments performed on 50 flies per genotype. One-way ANOVA with Dunnett's *post-hoc* test for multiple comparisons (**$p < 0.01$).

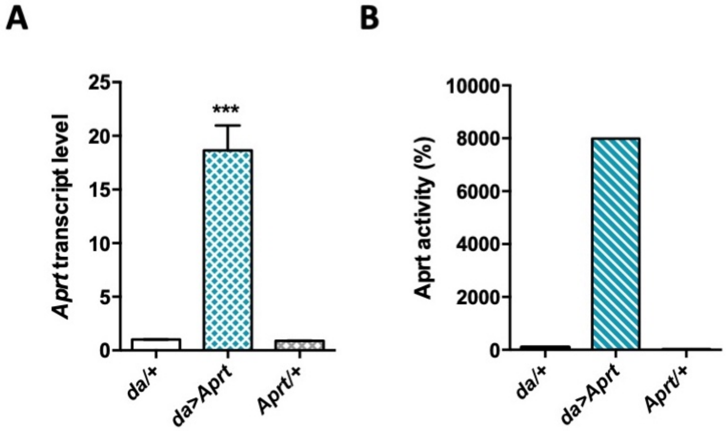


**Figure S7.** Transgenic expression of *Drosophila* Aprt. (**A**) A *UAS-Aprt* line was generated to allow for *Aprt* expression in specific cells. Upon ubiquitous expression with the *da-Gal4* driver (*da*>*Aprt*), *Aprt* mRNA level was found to be increased 18 times in adult heads compared to the driver (*da*/+) and effector (*Aprt*/+) controls. One-way ANOVA with Tukey's *post-hoc* test for multiple comparisons (***$p < 0.001$). (**B**) Aprt activity was also increased as much as 80 times in whole body extracts of the *da*>*Aprt* flies compared to the two controls. Results of one experiment performed on 20-30 male heads for RNA extraction and 20 whole flies for Aprt activity.

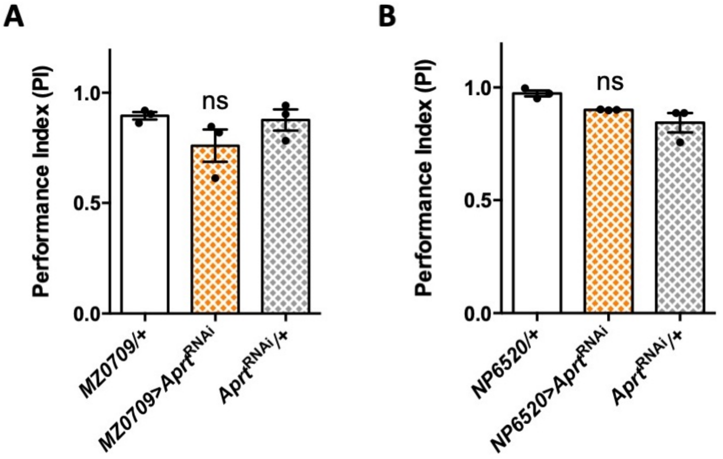


**Figure S8.** Downregulation of *Aprt* expression in the ensheathing glia does not alter locomotor performances. *Aprt* downregulation targeted to the ensheathing glial cells using *MZ0709-Gal4* (**A**) or *NP6520-Gal4* (**B**), did not impair startle-induced climbing of the flies at 10 d a.E. Results of 3 independent experiments performed on 50 flies per genotype, One-way ANOVA with Tukey's *post-hoc* test for multiple comparisons, ns: not significant.

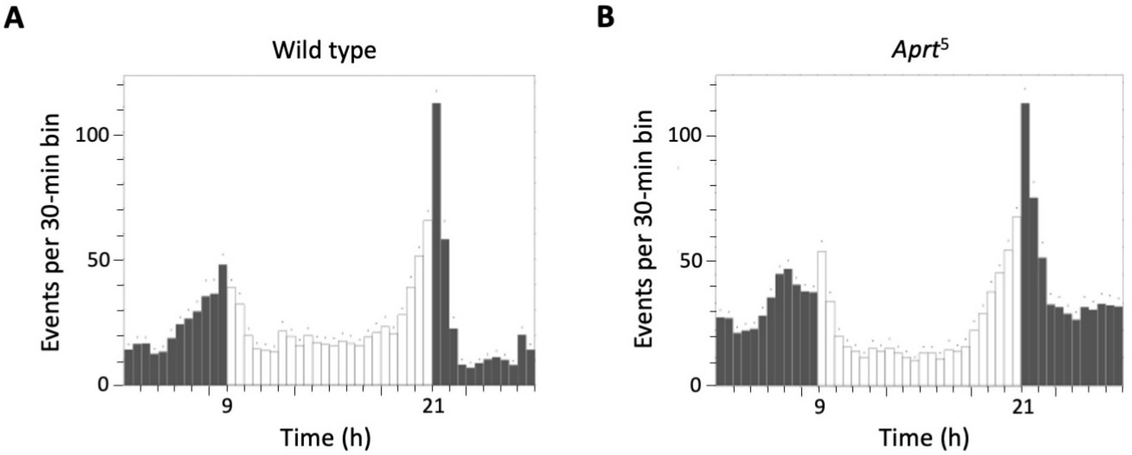


**Figure S9.** Daily locomotor activity profiles of wild-type and *Aprt*$^5$ mutant flies. Actograms showing activity profiles during a 12h:12h light-dark (LD) cycles from wild-type (**A**) and *Aprt*$^5$ (**B**) male flies at 8 d a.E. *Aprt* deficiency does not seem to alter circadian activity.

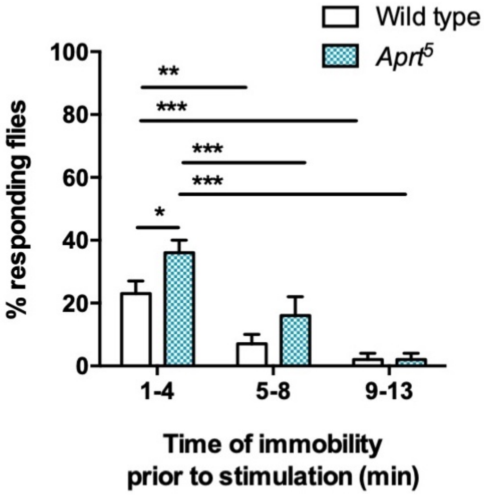


**Figure S10.** Wild-type and *Aprt*$^5$ mutant flies display similarly decreasing responses to stimulations following periods of immobility exceeding 5 min, confirming the standard criteria for sleep. Flies were exposed to a brief (1 sec duration) and mild vibration stimulus once every hour during two days. Mann-Whitney U test (*p < 0.05; **p < 0.01; ***p < 0.001).

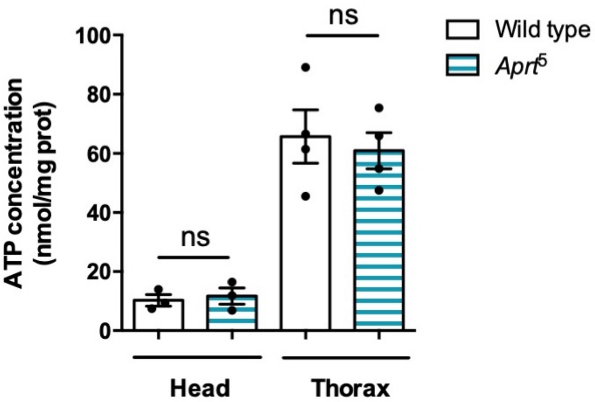


**Figure S11.** ATP levels are not altered in head and thorax of *Aprt*$^5$ flies compared to the wild-type. ATP was measured by a bioluminescence assay. Mean of 3 or 4 independent experiments performed on 30 heads or 5 thoraces per genotype. Unpaired Student's *t* test, ns: not significant.

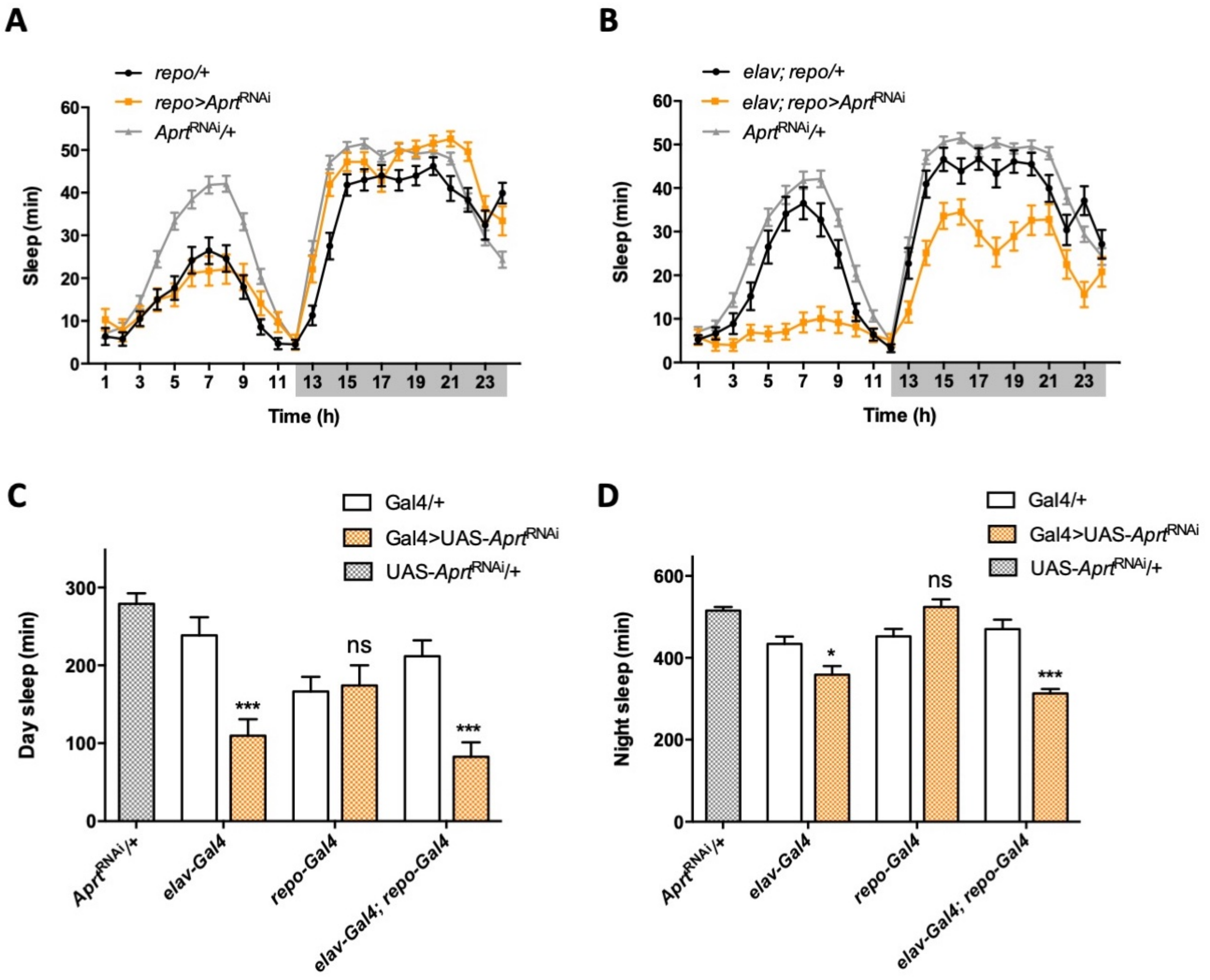


**Figure S12.** Sleep patterns of flies with cell-specific *Aprt* deficiency. (**A**) Sleep pattern of *repo>Aprt*$^{\mathrm{RNAi}}$ flies, showing that knock-down of Aprt in all glial cells did not induce any sleep defect. (**B**) In contrast, knocking down Aprt in all neurons and glial cells (*elav; repo>Aprt*$^{\mathrm{RNAi}}$) led to sleep reduction during both day and night, as with pan-neuronal *elav-Gal4* driver alone (shown in Figure 4F). (**C-D**) Quantification of sleep amount during day (**C**) and night (**D**) when *Aprt* was knocked down in all neurons (*elav-Gal4*), all glial cells (*repo-Gal4*) and both neurons and glial cells (*elav-Gal4*; *repo-Gal4*). Sleep reduction was observed when *Aprt* was downregulated in neurons but not in glial cells, and sleep was not significantly more affected when the double driver for neurons and glial cells was used, compared to the pan-neuronal driver alone. One-way ANOVA with Tukeys's *post-hoc* test for multiple comparisons (*$p < 0.05$; ***$p < 0.001$; ns: not significant).

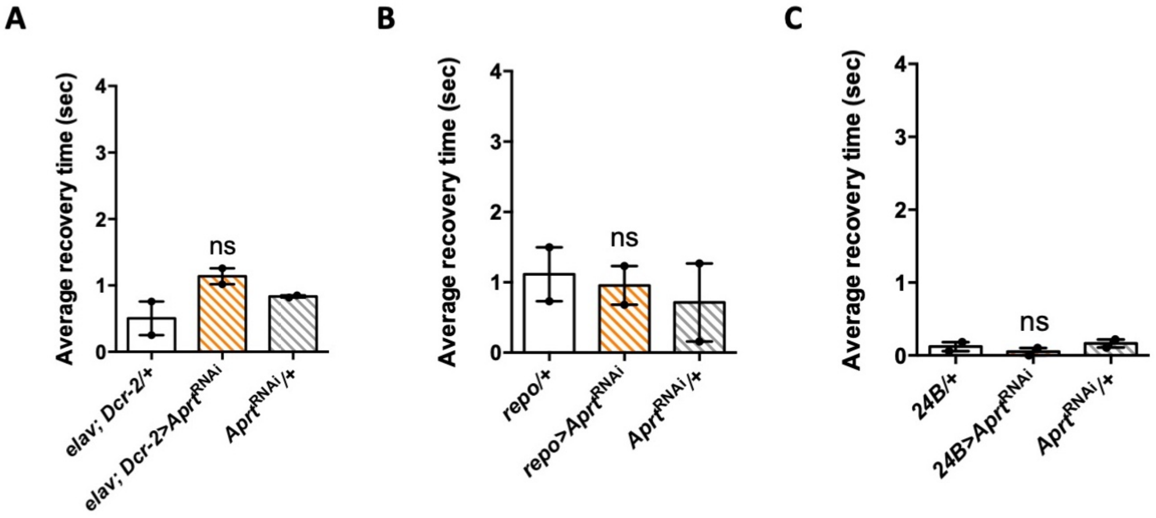


**Figure S13.** *Aprt* knockdown selectively in neurons, glia or muscle cells did not induce bang-sensitivity. (**A-C**) Downregulation of *Aprt* by RNAi either in all neurons (**A**), all glial cells (**B**) or all muscles (**C**) did not induce a seizure phenotype, in contrast to the effects of ubiquitous downregulation. Results of 2 independent experiments performed on 50 flies per genotype at 30 d a.E. One-way ANOVA with Tukeys's *post-hoc* test for multiple comparisons; ns: not significant.

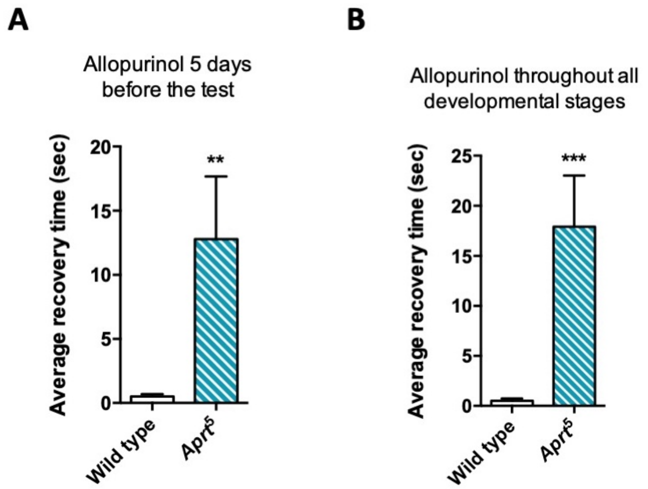


**Figure S14**. Administration of allopurinol does not rescue the bang-sensitivity phenotype of *Aprt*-deficient mutants. Feeding the *Aprt*$^5$ mutants with allopurinol at the same concentration used for uric acid normalization (100 µg/ml) either in adults 5 days before the test (**A**) or throughout all developmental stages (**B**) did not prevent the bang sensitivity of these flies. Results of one experiment performed on 50 flies per genotype at 30 d a. E. Unpaired Student's *t* test (***p* < 0.01; ****p* < 0.001).

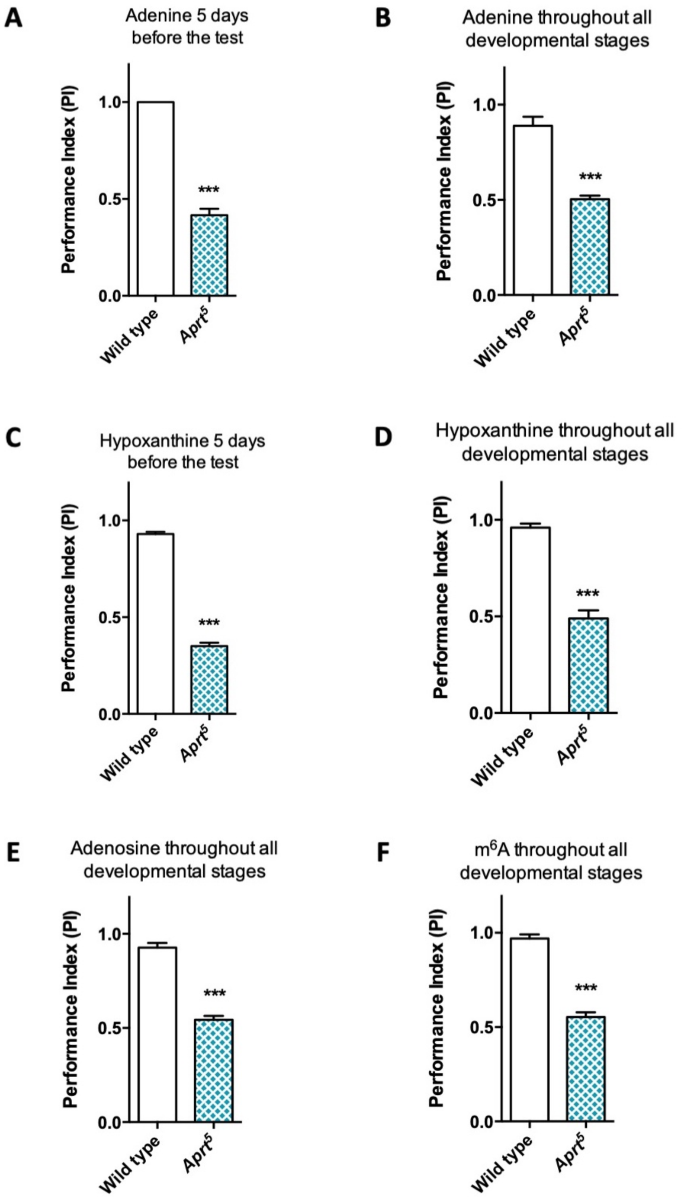


**Figure S15.** Administration of various purine compounds does not rescue the motricity defects of *Aprt*-deficient mutants. (**A-D**) Feeding the *Aprt*$^5$ mutant flies with adenine (**A**, **B**) or hypoxanthine (**C**, **D**) at 100 µM, either in adults 5 days before the test (**A, C**) or throughout all developmental stages plus 5 days before the test (**B, D**) did not rescue SING behavior defects. (**E, F**) Feeding the *Aprt*$^5$ mutants with adenosine (**E**) or $N^6$-methyladenosine (m$^6$A) (**F**) at 500 µM throughout all developmental stages plus 5 days before the test did not rescue SING behavior defects. Results of one experiment performed on 50 flies per genotype at 10 d a. E. Unpaired Student's *t* test (***$p <$ 0.001).

### Supplementary tables

**Table S1.** Aprt activity in wild-type and *Aprt*-deficient flies.

| Genotypes | Sex | Aprt activity (nmol/min/mg prot) |
|---|---|---|
| **Wild type** | males | 1.32 ± 0.17 |
| | females | 2.77 ± 0.27 |
| ***Aprt⁵/Aprt⁵*** | males and females | 0.04 ± 0.02 |
| ***Aprt⁵/Df(3L)ED4284*** | males | 0.02 ± 0.01 |
| ***da/+*** | males | 2.78 ± 0.41 |
| ***da>Aprt*** $^{RNAi}$ | males | 0.10 ± 0.01 |
| ***Aprt*** $^{RNAi}$ ***/+*** | males | 2.16 ± 0.37 |

Aprt enzymatic activity was assayed in whole adult extracts of wild-type Canton-S *Drosophila*. Females had more activity than males. No Aprt activity was detected in homozygous *Aprt*$^{5}$ or hemizygous *Aprt*$^{5}$/*Df(3L)ED4284* mutants. *Aprt* RNAi knockdown in all cells using *da-Gal4 (da>Aprt*$^{RNAi}$) resulted in an almost complete reduction in enzymatic activity of 96.4% and 95.4% compared to *da*/+ and *UAS-Aprt*$^{RNAi}$/+ controls, respectively. Results are mean ± SEM of 2-4 independent experiments performed on 20 whole flies per genotype.

**Table S2.** HGPRT activity in transgenic flies expressing wild-type or a pathogenic mutant form of human *HPRT1*.

| Genotypes | HGPRT activity (nmol/min/mg) |
|---|---|
| ***da/+*** | 0 |
| ***da>HPRT1-WT*** | 13.88 ± 3.75 |
| ***da>HPRT1-I42T*** | 2.70 ± 1.44 |
| ***HPRT1-WT/+*** | 0 |
| ***HPRT1-I42T/+*** | 0 |

Enzymatic assay performed on fly extracts revealed a significant amount of HGPRT activity in *da>HPRT1-WT*-flies and a much lower activity in *da>HPRT1-I42T* flies. No HGPRT activity was detected in *da*/+ *and UAS-HPRT1-WT*/+ or *UAS-HPRT1-I42T*/+ control lines, consistent with the lack of *HPRT1* homologue in the *Drosophila* genome.

**Table S3.** Primers used to generate *UAS-Aprt* and *UAS-HPRT1* constructs

| Genes | Primers | Sequences |
|---|---|---|
| ***Aprt*** | Ap-S1 | 5'-AGGGAATTGG**GAATTC**GTTATCAGTCGACATGAGCCC |
| ***Aprt*** | Ap-AS1 | 5'-ACAAAGATCC**TCTAGA**TCTAGAAAGCTTTCAGTACTTAATG |
| ***HPRT1*** | Hg-S1 | 5'- AGGGAATTGG**GAATTC**AAGAAGGAGATACAAAATGGC |
| ***HPRT1*** | Hg-AS1 | 5'- ACAAAGATCC**TCTAGA**GCTCGGATCCTTATCATTAC |
| ***HPRT1-I42T*** | Hg-I42T-S1 | 5'-CAGTCCTGTCCATA**G**TTAGTCCATGAGGAATAAACACCCT |
| ***HPRT1-I42T*** | Hg-I42T-AS1 | 5'-AGGGTGTTTATTCCTCATGGACTAA**C**TATGGACAGGACTG |

The added restriction sites are in bold type. The bases modified to match the *Drosophila* translation initiation consensus sequence are underlined. The mutation added is in red.

**Table S4.** Primers used for RT-qPCR experiments

| Genes | Primer sequences |
|---|---|
| ***AdoR*** | (forward) 5'-GGAGAAATTGCGATCGGATGACAC<br>(reverse) 5'-TCTTCAGCGAACTCCGAGTGAATG |
| ***Aprt*** | (forward) 5'-AATCAGCGCGGAAGACAAGCTA<br>(reverse) 5'-CCACCTTGCCGATGAGTTCAGT |
| ***DTH1*** | (forward) 5'-GGATCGAAAGCCAACCAAGTG<br>(reverse) 5'-CTTGGGGACCAACTGCGCTTTA |
| ***Ent2*** | (forward) 5'-ACGGCAAGGGATCAACGTC<br>(reverse) 5'-CCGTGCAGCAGGAATATAAAGA |
| ***rp49*** | (forward) 5'-GACGCTTCAAGGGACAGTATC<br>(reverse) 5'-AAACGCGGTTCTGCATGAG |

### Supplementary movies

**A** **B**

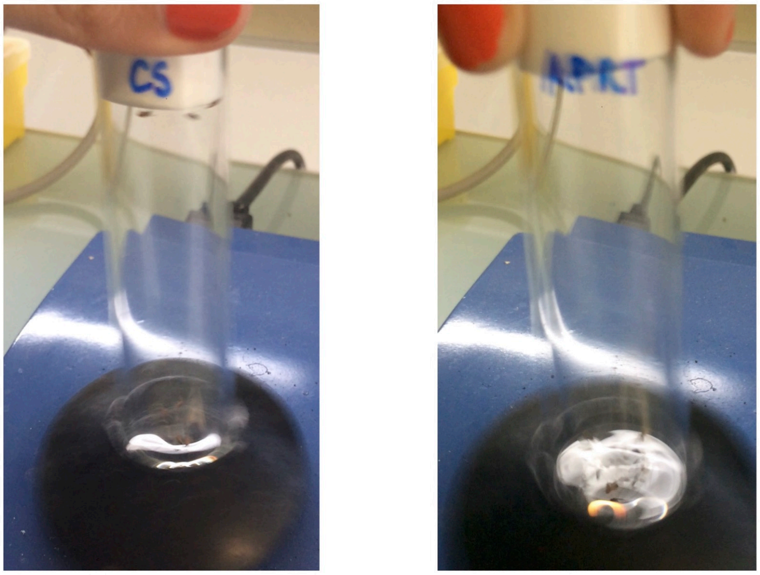


**Movie S1.** Bang-sensitivity phenotype of *Aprt*-deficient flies. (**A**) Wild-type Canton S flies (CS) are not very sensitive to 10-s vortexing and recover rapidly. (**B**) In contrast, *Aprt*$^{5}$ mutant flies (APRT) show seizure-like behavior and paralysis, and they recover slowly after such mechanical stimulation.
